# Cold Response of the Mediterranean fruit fly (Ceratitis capitata) in the Lab Diet

**DOI:** 10.1101/499228

**Authors:** Farhan J.M. Al-Behadili, Vineeta Bilgi, Junxi Li, Penghao Wang, Miyuki Taniguchi, Manjree Agarwal, Yonglin Ren, Wei Xu

## Abstract

Cold treatment at 0.0 °C with different exposure durations (0-12 days) was applied to the Mediterranean fruit fly *Ceratitis capitata* (Wiedemann) fed on lab diet. The examined developmental stages were early eggs (<6 hr) (E.E), late eggs (>42 hr)(L.E),first instar (1^st^), second instar(2^nd^) and third instar larvae (3^rd^). Pupation, adult emergence and sex ratios of survived flies were investigated to study the *C. capitata* responses to this low temperature treatment. Our results showed that based on pupation ratios, the 3^rd^ instar is the most cold tolerant stage with LT_99_=7.36 days. The second most cold tolerant stage is the 1^st^ instar with LT_99_=7.33 days. Cold tolerance at both two stages are very close, so they should be paid attention during the cold treatment. There were no significant differences on *C. capitata* sex ratios among different stages after treatment. This study improves our understanding of *C. capitata* responses to cold treatment, which may assist in the improvement of the current treatment strategies to control this destructive horticulture pest species.

## Introduction

The Mediterranean fruit fly, *Ceratitis capitata* (Wiedemann) (Diptera: Tephritidae) is one of the most destructive and invasive insect pests for horticulture biosecurity, global trade and world-wide phytosanitary [1]. *C. capitata* originated from sub-Saharan Africa and then spread throughout the Mediterranean region, Europe, the Middle East, Western Australia and the South and Central America [2].

*C. capitata* has been recorded feeding on over 300 fruit, vegetable and nut plant species. The main hosts include citrus, stone fruits, pome fruits, peppers, tomatoes and figs [3]. Plant hosts also include avocadoes, apricots, persimmons, strawberries, grapes, bananas, bitter melons, carambolas, coffees, guavas, peppers, papayas and blue berries. After mating, one female adult *C. capitata* can lay as many as 800 eggs in her lifetime. Since *C. capitata* attack happens when the majority of the production cost have already been expended, they cause huge horticultural industry losses. Therefore, *C. capitata* are regarded as one of the most destructive horticulture pest species. Furthermore, in recent years, the growing international trade of plant products increased the risk of introducing fruit flies across countries. Strict government policies were quickly made to mitigate these risks and minimize the damages. For example, pre-harvest actions including spraying, monitoring and inspections [4], with postharvest treatments such as fumigation, irradiation, heat or cold are required to control fly species [5]. Nevertheless, the current global trade of farm fresh products is suffering from the damages by various fruit fly species, prompting the need to develop more effective ways of biocontrol.

The available quarantine treatment technologies mainly consist of chemical (e.g. fumigation) and non-chemical treatments (e.g., cold, heat and irradiation). In recent years, with the discontinuation of several chemical products (e.g. Fenthion and Dimethoate), it has become even more urgent to rely on non-chemical postharvest control technologies to control fruit flies. Cold treatment is becoming an increasingly popular postharvest treatments due to the absence of chemical residues, mitigationes or mortality of the pest population; as well as increasing the strength of the fruits, and prolongings storage time [6–8]. It can also be applied to fruit at multiple stages, e.g. after packing and ‘in transit’ during lengthy transport by sea, as well as co-treatment with other postharvest treatments such as irradiation [9].

Optimal cold treatment conditions rely on the commodities, and most previous studies on *C. capitata* cold treatment reflect this, performing on the flies while they are within the fruits [6–8, 10–14]. On the other hand, because fruits vary in sizes, nutrients, compositions, and phytochemical profile, there were large differences in efficacy of heat transfer and fly development using similar cold treatment regime. For example, a previous study on the *C. capitata* cold tolerance in date and mandarin at 1.11 °C showed that *C capitata* is more sensitive to cold treatment when in date fruits than in mandarins [14]. Identification of the most cold tolerant development stage is crucial to determine the thoroughness of the treatment, however, there were also discrepancies on which *C. capitata* developmental stage was the most cold tolerant in previous reports. For example, Grout *et al.* [12] concluded that the most cold tolerant stage was the 2^nd^ instar based on a commodity group research report from South Africa. Hallman *et al.* showed that the 3^rd^ instar is the most cold tolerant stage [15]. We therefore find it compelling to conduct a research on cold tolerance of *C. capitata* reared on lab diet. The result will provide a fundamental baseline for comparison with data obtained from those conducted within fruits, and will be essential to establish cold response model, considering fruit sizes, compositions, nutrients and other variables. The overall objective of this study was to evaluate and understand *C. capitata* responses to low temperature. Based on these understandings, our long-term goal is to optimize current postharvest treatments and develop more environmentally-friendly, cost-effective, and efficient treatments for controlling *C. capitata*.

## Materials and Methods

### Insect Culture

*C. capitata* used in this study was originally established in 2015 from a laboratory colony maintained at the Department of Primary Industries and Regional Development’s (DPIRD), which has been periodically supplemented with the introduction of wild flies. Mature females lay eggs through the mesh (cloth sidewalls of the cages), which were collected and transported to the artificial rearing medium (Tanaka, Steiner, Ohinata, & Okamoto, 1969). After 13-16 days, pupae were collected and transferred into the adult breeding cages. The emerged adult flies were reared on the yeast hydrolysate, crystalline sugar and water. Rearing conditions were 26.0 ± 1.0 °C, 60 - 65% room humidity, and darkness light cycle of 16:8 hrs [11]. Early eggs (<6 hr), late eggs (>42 hr), 1^st^ instar, 2^nd^ instar and 3^rd^ instar larvae were used in this study.

### Cold Treatment Rooms

The cold room is a prefabricated unit with walls and ceilings of 100 mm expanded polystyrene. Joints and base of rooms are sealed with silicon sealant under aluminium covering extrusions. The floor is concrete; the door is 1500 mm wide × 1900 mm high × 100 mm thick with expanded polystyrene for insulation. The room dimensions are 4.33 m wide × 3.77 m length × 2.00 m high [11].

Refrigeration for the cold room is supplied by 2 × Patton (Model CCH 250) air cooled Condensing Unit with R22 refrigerant + 2 × Patton BL 38 Induced Draught Evaporator with refrigeration capacity of 5090 Watts at 0 °C. Temperature of the room is controlled through a surface mounted electronic thermostat (DIXEL, Italy) having a temperature range from −50 to +110 °C with a probe installed in the return air path. Up to 4 × Defrost cycles can occur per 24 hours if required. Two fans in the room (300 mm 5 blade propeller types) circulate air across the evaporator at an air flow rate of approximately 960 litres/second measured at various points in the room.

To maintain consistent temperature and to avoid temperature changes when opening the door of cold rooms, tested flies were placed in a chamber (1.2 m × 0.8 m × 0.6 m = 0.576 m3) located within the cold rooms. The temperatures inside the chamber were recorded every 30 minutes by Applent Multi-channel Temperature recorder (AT4508-128). The sensors were set inside the chamber in different levels. All sensors were calibrated in ice water (0 °C) before and after the treatment.

### Cold Treatment Tests

Carrot media was taken from a fridge approximately 24 hr before usage and was placed in sterile 90 mm plastic petri dishes at room temperature. The petri dishes were covered with lids and set aside on lab bench until infestation with insect of various stages. Approximately, 55 g of carrot media was added per petri dish with a clean spoon, making sure that the media was not aggregated or in contact with the lid. This ensured that larvae, if hatched, were able to move freely through the media and were not stuck to the lid. The lids from the control petri dishes were removed when larvae were first observed during incubation; the same number of days were then applied for treatment petri dishes when removing the lids.

Freshly laid *C. capitata* eggs were collected over a period of one hour prior to the trial and allowed to settle in a clean glass beaker containing 100 mL double distilled water. To count the number of eggs (in this study 100), droplets of water with suspending eggs were placed on a sterile petri dish using a transfer pipet. Eggs inside each droplet were counted using a manual cell counter, carefully picked up using a transfer pipet and placed onto the carrot media. Once the petri dishes were ready with the eggs, they were placed in a tray and transported to the cold rooms, where they were carefully placed in stacks of six replicate per exposure time inside the acrylic box.

100 eggs or larvae were collected, counted and transferred onto one petri dish with carrot diet (Table 1). Six replicates were prepared to be exposed from 0 to 12 days at 0.0 °C. After treatment, insects (six petri dishes each containing 100 individuals were retrieved at regular intervals of 24 h) were placed in a 750 mL clear disposable container containing a layer of sand and kept in an incubation chamber at 26.0 ± 1.0°C; 60 - 65% RH. Pupae emerging over a period of approximately 4 weeks were recorded. Untreated controls were kept in the same conditions (at 26.0 ± 1.0 °C; 60 - 65% RH) for pupae counting. Table 1 shows the total number of each *C. capitata* stage experimented in this study.

**Table 1.** The number of *C. capitata* used in each developmental stage.

| **Insect developmental stage** | **Insect numbers** | **Number of Replicates** | **Exposure time** | **Total Number of stages in each test** |
| --- | --- | --- | --- | --- |
| **Early eggs**<br>**(<6h)** | 100 | 6 | 0-12<br>days | 7200 + 600 control=7800 |
| **Late eggs**<br>**(>42h)** | 100 | 6 | 0-12<br>days | 7200 + 600 control=7800 |
| **1st instar** | 100 | 6 | 0-12<br>days | 7200 + 600 control=7800 |
| **2nd instar** | 100 | 6 | 0-12<br>days | 7200 + 600 control=7800 |
| **3rd instar** | 100 | 6 | 0-12<br>days | 7200 + 600 control=7800 |
| **Total** | 39000 insects (7800 × 5 bioassays) |  |  |  |

At the end of each exposure to cold treatment, the petri dishes were recovered and incubated at 26 °C; 65% RH as previously described. The petri dishes were carefully placed on a thin layer of clean sand in a 750 mL rectangular disposable container. A piece of mesh cloth has been put on top of the container and the lid was then affixed. The lids were prepared to have six holes ensuring air circulation when placed in the incubator. Control petri dishes were checked regularly and on day 8 when instars were visible, the lid was removed and the remaining configuration was left as described. The lids of the petri dishes in treatment groups were removed after being placed in the incubator as follows: after 7-8 days for early eggs trial; after 5-6 days for late eggs; after 3-4 days for first instar larvae; after 1-2 days for second instar larvae. Third instar larvae had no lids from the start. The experiment set-up, number of eggs/larvae and treatment periods were shown in Table 1. During each trial, the temperature were recorded at 30-minute intervals in each cold room, at the carrot diet, and in the air of the chamber. The sand was renewed three times over six weeks to collect pupae. Pupae, emerged adults and the sex ratios were recorded and analyzed.

To study the effects of cold treatment on eggs, mortality was recorded by comparing a total number of pupae or adults produced, and eggs that were incubated at 26.0°C were used as a control. This comparison provided mortality and eggs were claimed “live” if pupae or adults were produced after treatment. This procedure included sieving through sand containing pupae followed by counting the total number of pupae. Sieving was carried out using a metal mesh tray (1.6 mm) that allowed sand particles to pass through but retained pupae. Sieving was carried out as soon as pupae were first seen; with this process repeated three times until there were no more pupae found in the sand. Pupae were carefully placed on a clean surface, and any large sand particles that remained were separated using a glass slide. Pupae were then counted and placed in a sterile petri dish which sat on a laboratory bench at room temperature. Additional observations on the number of emerged flies were considered. This provided information on the viability of pupae and whether they were able to develop into adults. Three observations were made on the ratio of male: female. The mortality rate was determined by the number of viable pupae (or adults) recovered from the treated insects, relative to the total numbers treated. These values were then used for further analysis.

## Statistical analysis

The mortality rate of the insect under cold treatment was statistically estimated following the Median lethal time method (LT). The 90% and 99% (LT90 and LT99) were estimated. We have evaluated four different models separately for pupils and adults on every stage, including eggs, 1^st^ instar, 2^nd^ instar, and 3^rd^ instar. The best fitting model was selected for estimating the LT90 and LT99. The LT estimated under generalized linear model with both a probit and a logit link function on cold treatment days, which are the mostly used dose-response model where in our scenario days of treatment was considered as dose. The model can be written as:

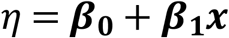

Where η is the response or proportion mortality, x is the dose, β0 is the intercept, and β1 is the coefficient of the dose. The evaluated 4 models include (1) probit model on log transformed treatment days; (2) logit model on lgo transformed treatment days; (3) probit model on treatment days without log transformation; and (4) direct logit model on treatment days. The best model was selected based on three different criteria, including exploratory analysis, BIC value, and regression residues. The best fitting model following these criteria for each insect stage was finally selected to estimate LT90 and LT99. This demonstrates that we are using the best fitting model for LC estimation. The 95% confidence interval was also reported. R statistical environment (version 3.3.2) with base library was used to estimate the LT and confidence interval, ggplot2 package was used for generating plots.

## Results

In this project, a total of ~39,000 *C. capitata* eggs or larvae were used for cold treatment. 7,800 eggs/larvae were used for each stage (early eggs, late eggs, 1^st^ instar, 2nd instar and 3^rd^ instar larvae). The duration of treatment ranged from 0 to 12 days at 0.0 °C to reach 100% mortality (Table 2). During the cold treatment, probes were used to monitor the temperature in the cold room, and the results showed that the temperature was stable at 0.0±0.2 °C, from the start to the end of the experiment.

The effects of cold treatment on *C. capitata* were shown in Table 2 and Fig. 1. Pupation and adult emergence ratios were used to calculate the mortality rates. By using pupation, if an egg/larvae could not develop to pupae after treatment, it was defined “death”. Similarly, by using adult emergence, if an egg/larvae could not develop to an adult after treatment, it was defined as “death”. Our results showed that the five developmental stages (early egg, late egg, 1^st^ instar, 2^nd^ instar and 3^rd^ instar) differed in their cold tolerance at 0.0 °C. By using the recovered pupation (Table 2 and **S1** Fig.), 1st instar larvae is the most cold-tolerate stage as it took nine days at 0.0 °C in the lab diet to reach zero pupae (100% moratlity). 3^rd^ instar is the second most cold-tolerate stage, which took seven days at 0.0 °C to reach 0 pupae. Early eggs, late eggs and 2^nd^ instar need six days to reach zero pupae after treatment. Interestingly, using the emerged adults to study the mortality achieved slightly different results (Table 2). For example, early eggs and 2^nd^ instar all need six days to achieve zero adults (100% mortality) from 600 eggs/larvae. Late eggs and 3^rd^ instar both need five days to achieve zero emerged adult, thus suggesting late eggs have a greater susceptibility to cold than early eggs. 1^st^ instar larvae is the most cold-tolerate stage as it took the most days (9 days) to reach 100% moratlity.

**Figure 1.**
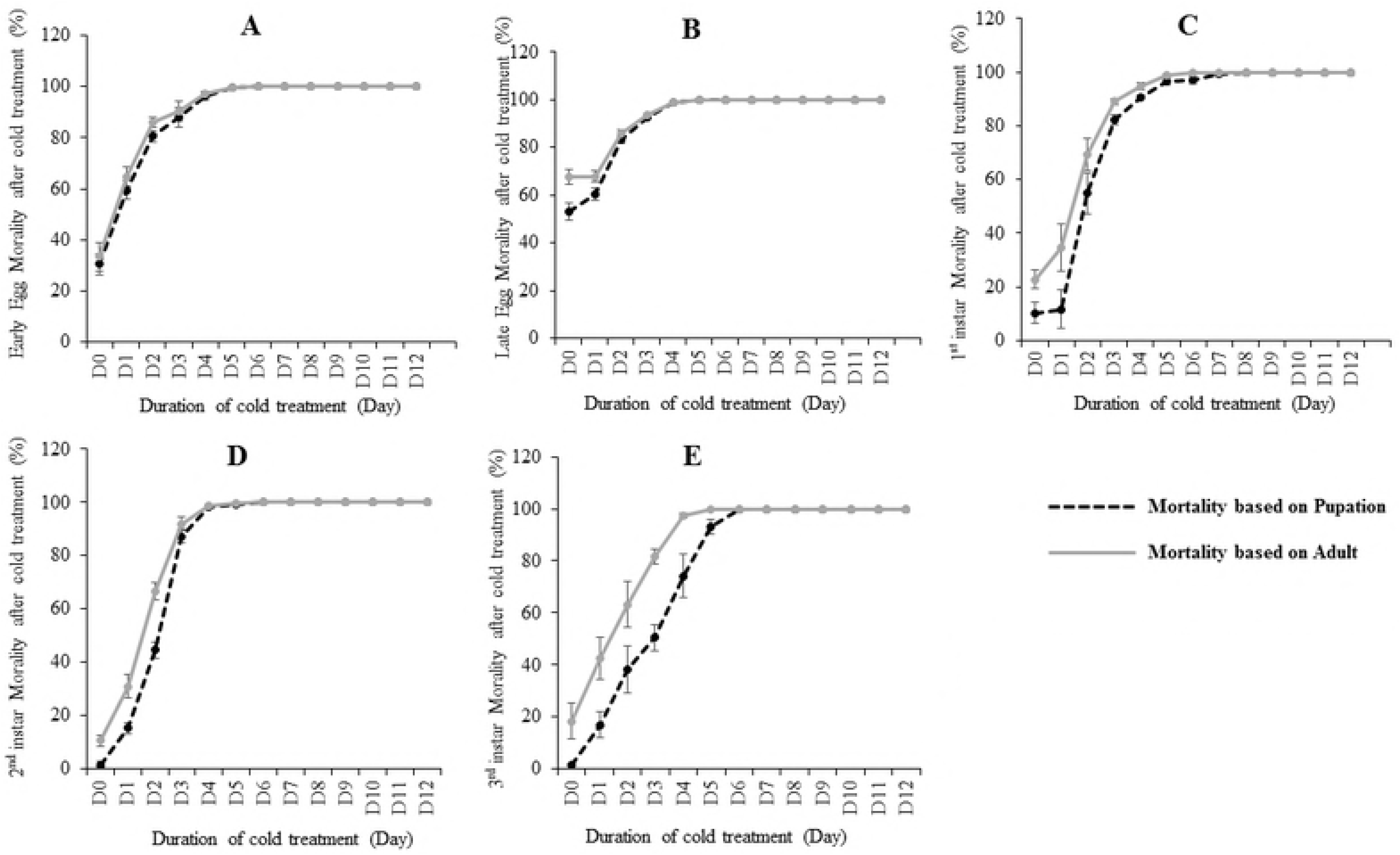
The mortality (%) calculated based on pupation and adults at fife bioassay. Error bar means standard error.

The pupation or adult emergence ratios from the non-treated (control) eggs/larvae varied (**S1** Fig. A and B). The results for larvae stages (1^st^, 2^nd^ and 3^rd^ instar) are very similar (89-99% pupation and 77-89% adult emergence). The early eggs in the control showed 69.5% pupation and 66.6% adult emergence while the late eggs showed only 46.8% pupation and 32.3% adult emergence. The control eggs (including both early and late eggs) showed much lower pupation/adult emergence ratios than the control larvae (1^st^, 2^nd^ and 3^rd^ instar), which may be due to the low hatching ratios from eggs to larvae (Table 2).

As mentioned, four different models were selected separately for pupations and emerged adults on every stage and the best fitting model was selected for estimating the LC90 and LC99 (S1 Table). The four models include: log on days with probit, log on days with logit, no log on days with probit, and no log on days with logit (S1 Table). The results showed that no log models are better than log models in this study (S1 Table). When using pupation as the end point for mortality analysis, no log on days with probit model was selected for early egg, late egg and 3rd instar while no log on days with logit was selected for 1st and 2nd instar larvae. When using emerged adult as the end point for mortality analysis, no log on days with probit model was selected for late egg while no log on days with logit was selected for the rest stages. We modelled the duration of cold treatment to induce 90 and 99% mortality at five immature stages of C. capitata fed on a lab diet (Table 3). Using the recovered pupation as end point for mortality modeling, the results showed that the 3rd instar is the most cold tolerant stage with LT99=7.36 days, which is very close to the result from a previous study with LT99=7.0 days [15]. The second most cold tolerant stage is the 1st instar with LT99=7.33, based on pupation. Interestingly, based on the emerged adult ratios, LT99 was 5.71, 5.7, 6.4, 5.41, and 6.14 days for eggs, Late eggs, 1st instar, 2nd instar and 3rd instar respectively. By using loss of emerged adults as end point, our bioassay showed that 1st instar is the most cold tolerant (LT99=6.4) and 3rd instar is the second most cold tolerant stage (LT99=6.14). However, by using loss of pupae as end point, the modelling analysis results showed 3rd instar is the most cold tolerant stage (LT99=7.36) and 1st instar is the second most cold tolerant stage (LT99=7.33).

It was reported that in reptiles, the temperature of the eggs during a certain period of development is the deciding factor in determining sex. Therefore, to study if the cold treatment affects the sex development of the emerged adults, we compared the sex ratios of the treated eggs and larvae. First, we collected all the survival adults after treatments from five developmental stages and calculated the percentage of female. The results (Fig. 2A) showed that there are no significant differences at each stage, which are all close to 50%. Second, to examine if a certain period of days treatment influences the sex development, we analysed the female ratios from treated flies at different days (Fig. 2B). If the numbers of emerged adults are too low (<10), the adult sex ratios may be biased, which will not be included in this analysis. From Day 0 to Day 3, based on ANOVA test there are no significant differences on each day treatment of five immature stages (Fig. 2B). Overall, cold treatment at 0.0 degree C on C. capitata does not affect sex ratio.

**Figure 2.**
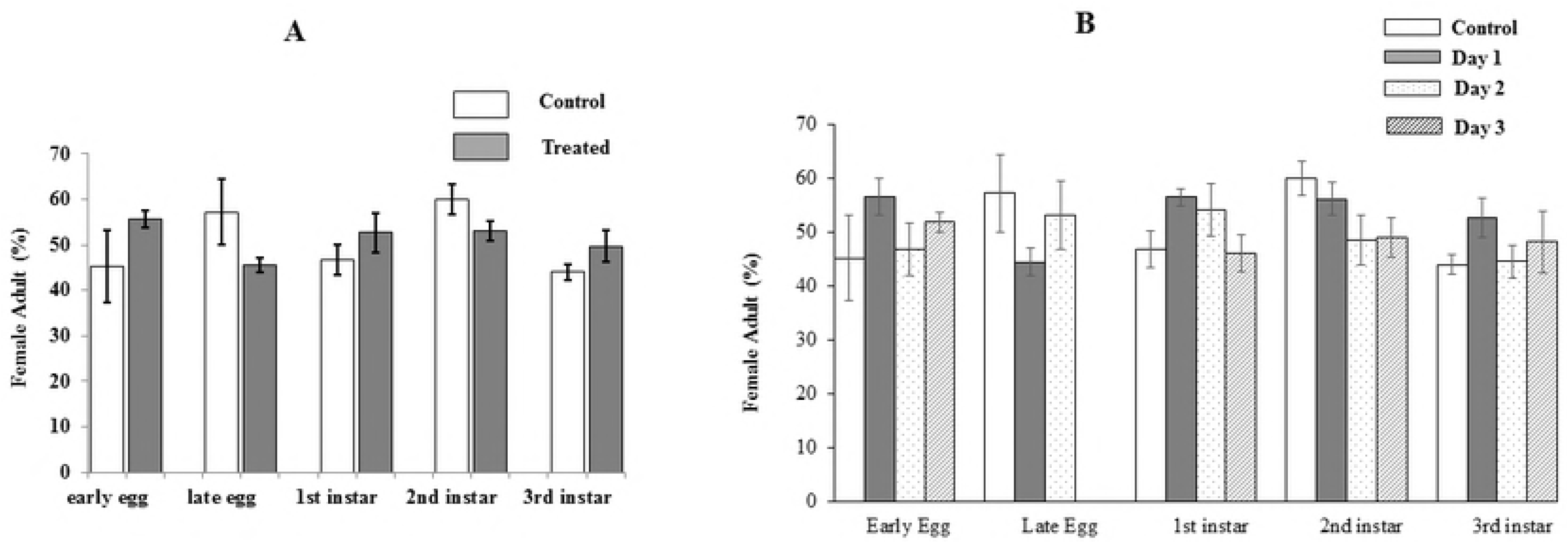
The average sex ratios during the first three days of treatment (A) and on each day (B). Error bar means standard error.

## Discussion

Cold treatment is a common method for eradicating fruit flies in fresh fruits and other products. It has been studied, analysed and incorporated into quarantine regulations [6, 8, 10, 11] and overseas countries [7, 12–14].

In this study, we chose to study *C. capitata* cold response from the lab diet but not fruits due to the variation in sizes, materials, nutrients and chemicals, resulting in significant differences in fly physiology and development. Secondly, some fruits may not be favorable hosts for fruit flies and so might be associated with the production of weak flies, resulting in the differences in the mortality rates. Furthermore, there are different infestation methods for various fruits. For example, by using the natural infestation, how many eggs were laid into a single fruit is hard to clarify. By using artificial infestation (e.g., injecting eggs or larvae into fruits), methods may affect fruits quality; fruit fly development and even cause bacterial or fungi contamination. All these can result in very different results in cold treatment experiments. Therefore, here we chose to use lab diets. As the high homogeneity of the diet in terms of ingredients, quantity, size and dimensions of experimental units make the results more reliable and reducing of experimental error. Because through this way, we can be clear about how many eggs/larvae were used in the analysis and avoid the fruit/infestation issues. Cold treatment analysis of *C. capitata* on lab diet have been performed [15], but not in a comparative way on all five different stages as we did here.

To evaluate cold treatment effects on *C. capitata*, the first question is how to define mortality. Eggs or larvae may survive from the cold treatment, but fail in pupation or adult emergence due to unknown effects. On the other hand, some of them succeed in adult emergence but fail in sex development and reproduction, so they are not biologically “alive” flies. Therefore, here we used the recovered pupae or adults as standards to help define our mortality here. If one egg/larvae fail in developing to pupae or adult after treatment, it is a “dead” fly.

Our bioassays (Fig. 1 and Table 2) showed that the 1^st^ instar is the most cold tolerate stage. However, modelling analysis LT99 results showed the 3^rd^ instar is the most cold tolerate based on pupation result and 1^st^ instar is very close to 3rd instar. Based on the emerged adults, LT99 results showed that 1^st^ in star is the most cold tolerant and 3^rd^ instar is the second. All these results suggest both 1^st^ instar and 3^rd^ instar are among the most cold tolerant stages, on which more attention should be paid in our postharvest treatment. The previous studies are not consistent concerning the most cold tolerant stage of *C. capitata*. There are several major reviews and seven annexes of International Standards for Phytosanitary Measures (ISPM) that deal specifically with the cold tolerance of this species. Grout et al. [12] concluded that the most cold tolerant stage was the 2^nd^ instar based on a commodity group research report from South Africa. Hallman *et al.* showed that the 3^rd^ instar is the most cold tolerant stage [15]. A study compared tolerance of eggs, a mixture of 1^st^ and 2^nd^ instars, and mostly 3^rd^ instars to 1.5±0.5 °C in oranges and found both larval groups to be very similar, with the younger instars showing a very slight advantage in survival [16]. Another study found that the 2^nd^ instar was the most tolerant to 1.0±0.2 °C in two cultivars of lemon [8]. The third study found that the 2^nd^ instar was more tolerant than the 3^rd^ in five types of citrus fruits at 2 and 3 °C [17]. There are plentiful differences in the experiment set up, including, but not limited to fruits, infestation methods, temperatures, fruit sizes, materials and nutrient. Therefore, it is not surprising that the detected most cold tolerate stage is different. It is also likely that different population of fruit flies vary in relative tolerance of the different states to cold treatment. It was reported previously that geographically isolated populations of Medfly differ in reproductive patterns, survival, developmental rates, intrinsic rates of increase [18]; suggesting their different tolerance to low temperatures.

In this study, we compared early eggs (< 6 hrs) and late eggs (>42 hrs) in the cold treatment because the embryo development stage may affect their biology, physiology and cold tolerance. For the fruit picked up and transferred to the cold storage, it is likely to collect some early eggs which were laid just before the fruit picking and late eggs which have been laid for a period of time. Egg is a fast developing stage so the age difference can be a significant issue that has been ignored in previous research. Here our results showed that the early eggs is a bit more tolerant than late eggs. We also compared the sex ratios of survived adults after treatments. Our result showed that there is no significant difference in the sex ratios at different stages.

Whilst a number of studies investigate industry-relevant applications of cold treatment of Tephritid fruit [5–7, 12] in various fruits at various stages, a fundamental understanding of the mechanisms underlying why and how cold can kill the flies is currently lacking. Without this knowledge, it is difficult for us to improve and optimize our current cold treatment strategies and develop more efficient treatment methods. A number of technology breakthroughs in genomic research, such as low-cost RNA sequencing (RNASeq) libraries, high-throughput quantitative polymerase chain reaction (qPCR), bioinformatic software and gene network reconstruction, have enabled advances to be made in biomedical and livestock science. With the recent publication of the genome of *C. capitata* [19], there is now an attractive opportunity to apply these technologies to *C. capitata*. This will underpin a real breakthrough in understanding stress-based fruit fly infestation, which has become even more important.

## Supporting information

**S1Table. Four different models were used in the C. capitate mortality data analysis.**

**S1 Fig. (A) Mortality (%) based on pupation and adults (B) ratios from the treated *C. capitata*.**

## Acknowledgments

We would like to thank National Centre for Post-harvest Disinfestation Research on Mediterranean Fruit Fly in Murdoch University, Dr Sonya Broughton and Emma Mansfield from Department of Primary Industries and Regional Development (DPIRD) for the help in this project. We also thank Dr Yalin Liao for proof reading. Dr Wei Xu is the recipient of an Australian Research Council Discovery Early Career Researcher Award (DECRA) (DE160100382).

